# Detection of *GBA* missense mutations and other variants using the Oxford Nanopore MinION

**DOI:** 10.1101/288068

**Authors:** Melissa Leija-Salazar, Fritz J. Sedlazeck, Katya Mokretar, Stephen Mullin, Marco Toffoli, Maria Athanasopoulou, Aimee Donald, Reena Sharma, Derralynn Hughes, Anthony H Schapira, Christos Proukakis

**Affiliations:** Department of Clinical Neuroscience, Royal Free Campus, Institute of Neurology, University College London, London, United Kingdom; Human Genome Sequencing Center, Baylor College of Medicine, Houston, USA; Institute of Translational and Stratified Medicine, Plymouth University Peninsula School of Medicine.; Department of Molecular Neuroscience, Institute of Neurology, University College London, London, United Kingdom; Department of Paediatrics, Royal Manchester Children’s Hospital; The Mark Holland Metabolic Unit, Salford Royal Foundation NHS Trust, Salford, UK; Institute of Immunity and Transplantation, Lysosomal Storage Disorders Unit, Royal Free Hospital, London, UK

**Keywords:** GBA, Parkinson’s disease, Gaucher disease, Long-read sequencing, Oxford Nanopore MinION, mutation detection, mutation phasing

## Abstract

Mutations in *GBA* cause Gaucher disease when biallelic, and are strong risk factors for Parkinson’s disease when heterozygous. *GBA* analysis is complicated by the nearby pseudogene. We present a simple and efficient method to sequence *GBA* as an 8.9 kb amplicon on the Oxford Nanopore MinION. We use existing software for a rapid and comprehensive analysis, including blinded detection of all known missense mutations in our samples, with zygosity assessment and full phasing, as well as detection of intronic SNPs, and of an exonic deletion. Our protocol provides an efficient research and clinical diagnostic tool in this difficult gene.

## Background

The *GBA* gene encodes the lysosomal enzyme Glucocerobrosidase, deficiency of which leads to accumulation of glucosylceramide. Biallelic (homozygous or compound heterozygous) mutations in *GBA* cause Gaucher disease (GD), the most common lysosomal storage disorder.[1] Heterozygous *GBA* mutations are a significant risk factor for Parkinson’s disease (PD),[2, 3] with evidence of longitudinal changes in many carriers suggestive of prodromal PD.[4] *GBA* mutations are also associated with Dementia with Lewy bodies [5] and Multiple System Atrophy, [6, 7] related conditions which also demonstrate aggregation of the alpha-synuclein protein. At present, more than 300 mutations have been linked to Gaucher disease,[8] and the number of studies analysing the prevalence and phenotype of *GBA* mutations in PD is rapidly increasing.[9] [10, 11] [12, 13]

*GBA* comprises eleven exons and ten introns over ~8 kb on chromosome 1q21. A nearby pseudogene *GBAP* has 96% exonic sequence homology to the *GBA* coding region. The region also contains the Metaxin gene (*MTX1*), and its pseudogene. The existence of these two pseudogenes confers an increased risk for recombination between homologous regions, which can generate complex alleles. The homology between *GBA* and *GBAP* is highest between exons 8-11, where most of the pathogenic mutations have been reported, usually resulting from recombination events.[8]

The complex regional genomic structure complicates PCR and DNA sequencing, and some exons are also problematic in exome sequencing[14] and whole genome sequencing.[15] Established analysis protocols usually involve PCR of up to three fragments, carefully designed to not amplify *GBAP*,[16] followed by Sanger sequencing of coding exons. Illumina targeted sequencing protocols have also recently been developed.[9, 12] In recent years, long reads produced by sequencing DNA molecules in real time have become commercially available, and have several advantages over short reads.[17] Oxford Nanopore sequencing technology analyses a single DNA molecule while it passes through a pore, producing characteristic changes in current depending on the sequence.[18] The Oxford Nanopore MinION is currently the most portable long-read sequencer. It can be plugged into a computer through a USB connection and provides sequencing data and run metrics data in real time. It has been used for applications ranging from pathogen sequencing in the field,[19] to sequencing a whole human genome.[20] It is still not routinely used in human disease diagnostics, but has been successfully used for SNV detection *in CYP2D6*, *HLA-A* and *HLA-B*,[21] *TP53* in cancer [22] and *BCR-ABL1* in leukemia,[23] and for chromosome 20 in a recent whole genome sequencing study. [20]

In the present study, we present an efficient laboratory and bioinformatic protocol for *GBA* analysis using the MinION. In addition to disease-causing variants, it can detect intronic ones, and provide phasing information. The MinION protocol can thus provide further insights into *GBA* than other sequencing technologies, and is ready to be considered for diagnostic use.

## Results

### Validation of correct alignment and preliminary mutation detection

To test our method, we first performed sequencing with 2D reads on brain DNA, acquired over five runs, using two earlier chemistry versions (R7.3 and R9), and the Graphmap aligner. The basic metrics of all are shown in Tables S3- S4. Nine samples had *GBA* coverage >60 over these runs, and were analysed further. We confirmed that reads mapped to the *GBA* target region, with only ~1.1% of reads aligning to pseudogene (Table S4). Two of these samples were known to have heterozygous *GBA* coding variants (three SNVs comprising the “RecNciI” allele in S5, and p.L483P, or L444P in S8). These were called by Nanopolish, and were clearly visible on IGV (Figure 1). One additional coding SNV was detected in S5 (g.155205518C>G; rs1064651; p.D448H, or D409H). This can occur in *cis* with the RecNciI allele.[24] There were non-coding variants in all samples. One apparently novel intronic variant in sample S1 (chr1:g.155207565C>T; intron 6: c.762-196G>A) was confirmed by Sanger sequencing (Figure S2). It has now been reported as a very rare SNP in dbSNP build 150 (rs979955939; minor allele frequency 0.0001) and gnomAD genomes (1 of 30,762 alleles). Several other non-coding SNPs were detected (Table S4). One additional candidate, which was not a known SNP, was detected in S3. Review on IGV revealed that the same base change was present in all eight other samples, with AF 10-20% in five of them (Figure S3). NanoOK showed that A>G is the second commonest erroneous substitution in this sample (15.94%). Sanger sequencing did not confirm this variant, demonstrating that a common base error, with frequent reads supporting the same variant in most samples, is a false positive.

**Figure 1.**
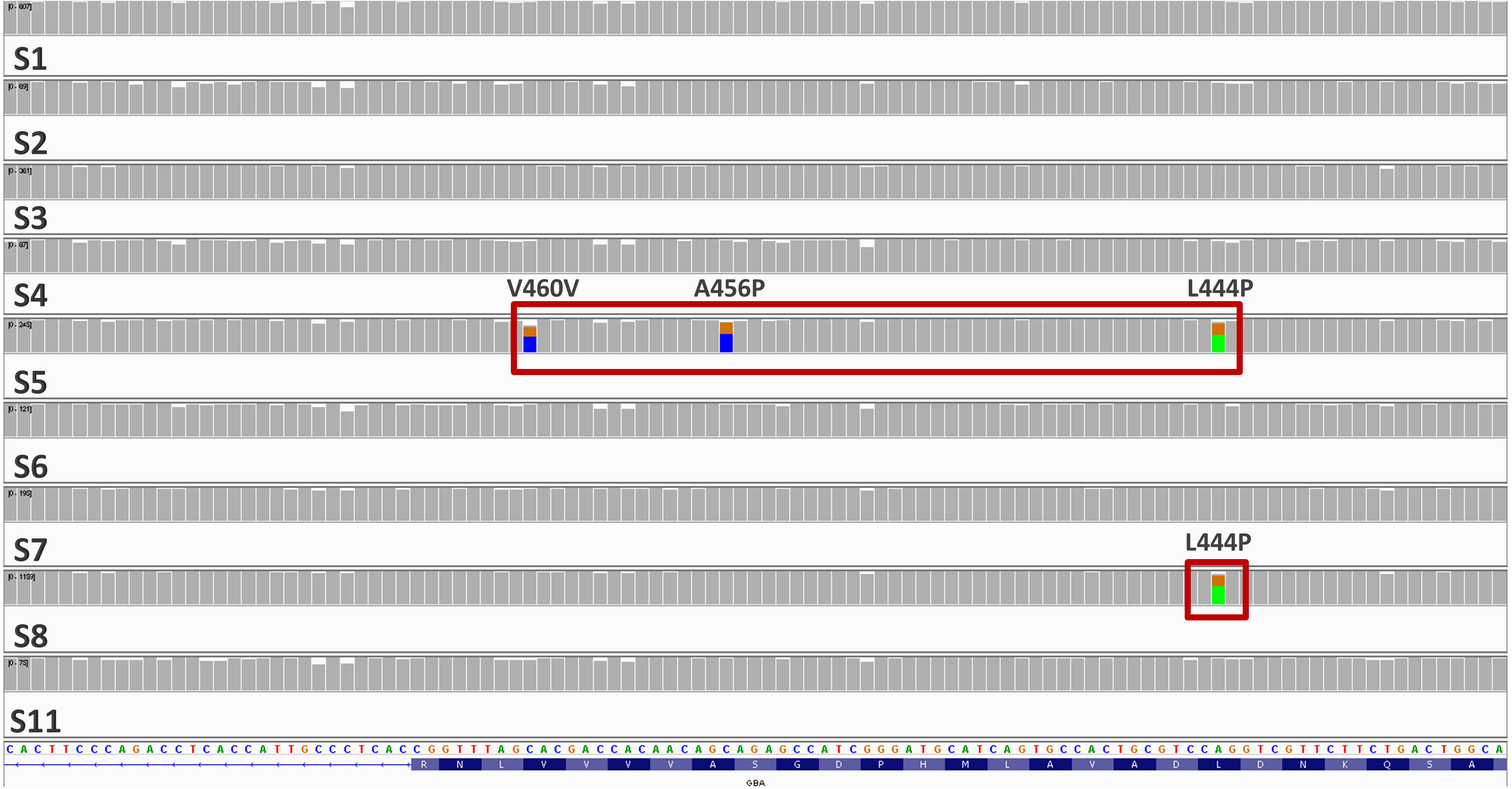
Detection of known variants in S5 and S8 in the validation phase. The IGV trace over exon 10 is shown for all samples sequenced with 2D reads.

### Use of newer chemistry to test detection in Gaucher patients

For the next part of the study, given the rapid improvements in Nanopore chemistry, and availability of the newer R9.4 cells, we decided to test samples known to carry pathogenic mutations, to determine the potential for diagnostic use. We used the Kapa PCR protocol, because of a possible minimal error reduction (Table S4). We included DNA from the two previously tested PD brain samples carrying RecNciI and p.L483P (S5 and S8), two additional untested PD cases (one brain and one saliva), and six saliva samples which had previously been determined to carry heterozygous or biallelic mutations. These comprised four GD patients and two carriers, although their status and previously established genotypes were not revealed until after the analysis was performed. We multiplexed these 10 samples on a R9.4 flow cell.

NanoOK analysis showed high base accuracy for all samples (mean 93.2%) (Tables S3 and S5). We aligned data using both Graphmap, and the newly developed NGMLR, with a mean *GBA* coverage >300, and minimal number of reads aligning to the pseudogene (average 0.78% and 1.97% of the reads aligning to gene with Graphmap and NGMLR respectively; Table S5).

### Coding mutations are detected

We called variants using Nanopolish (version 0.8.4) on data aligned both with GraphMap and NGMLR. We first focused on coding SNVs, which were detected in eight of the ten samples, regardless of the aligner used (Table 1; Figure 2). The two untested PD patients S16 and S18 were negative. We detected all previously known coding missense mutations, at the correct zygosity. These included p.N409S (N370S) in three GD cases, in the homozygous state in two (S12, S14), and heterozygous in one (S17) (Figure 2A), and the second mutation in S17 (p.L105P; Figure 2B). In another GD patient we detected two other heterozygous pathogenic mutations (p.R502C, p.R535C; Figure 2C,D). In the “RecNciI” carrier (S5), the additional p.D448H mutation was confirmed (Figure 2F), which would lead to this allele being designated as “RecTL”. We also detected heterozygosity in three samples from individuals without GD for p.L483P (L444P) (Figure 2E), including the one tested earlier. The mean quality score for coding heterozygous SNVs was 638 (standard deviation 229), and the lowest 337.8. The lowest scores were in S17, which had the second lowest coverage (205.1). There was a non-significant trend for coding heterozygote SNVs in a sample to have higher mean quality scores with higher coverage (Spearman correlation r=0.77, p=0.10, for the six samples which had at least one). The cut-off for a true positive may therefore partly depend on coverage.

**Table 1.**
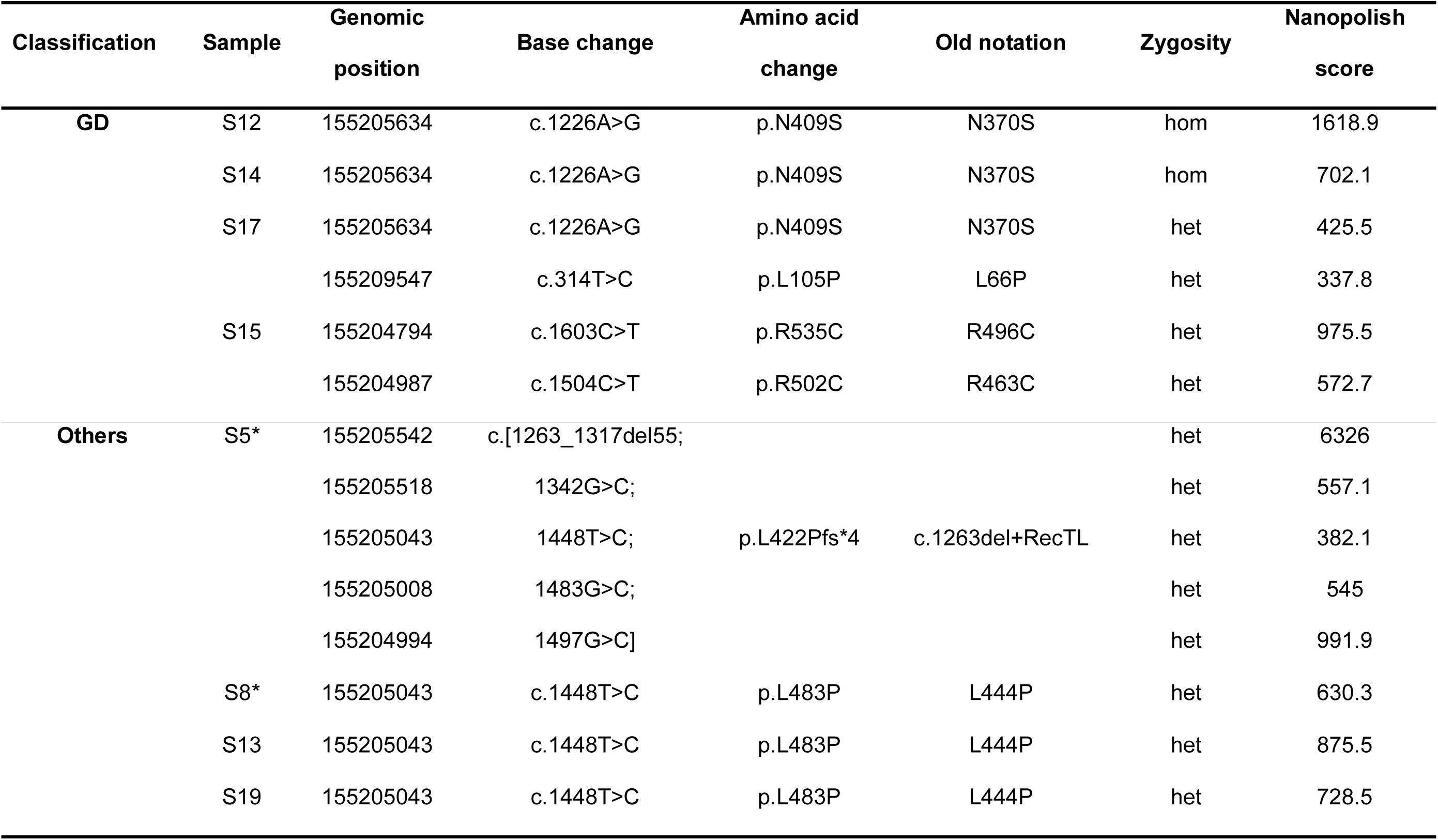
Coding mutations detected. The samples were classified as GD (where two mutations were expected), and others (where up to one was expected). * indicates sample included in early 2D read work. The old aminoacid notation is included. Zygosity is shown for each mutation in each sample (het=heterozygous, hom=homozygous). The Nanopolish quality score is shown for the NGMLR aligned data.

**Figure 2.**
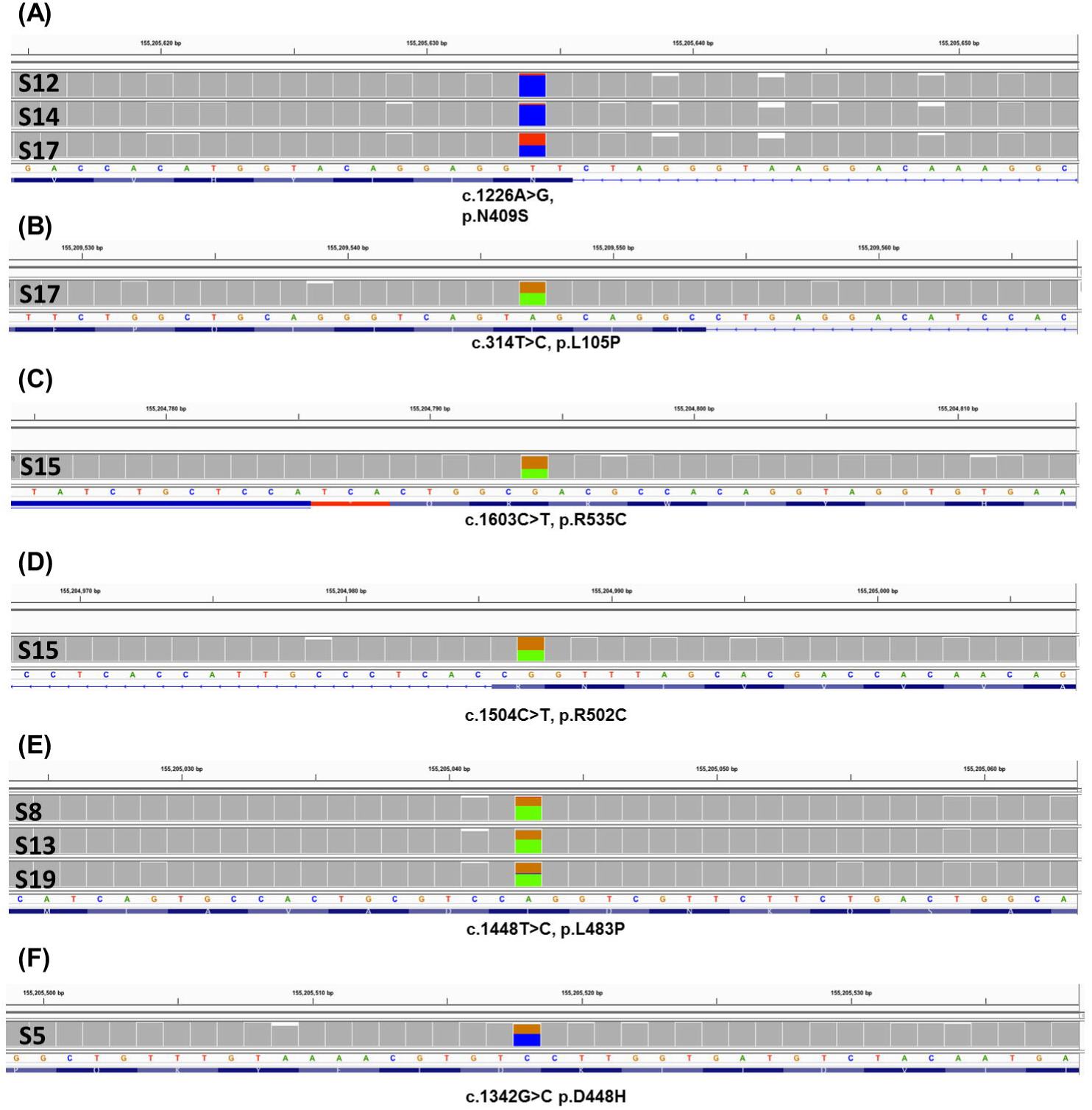
Missense mutations detected with R9.4 chemistry. The IGV trace is shown for each sample with a mutation. The mutated base is shown, with 20 bases on either side. The three SNPs which comprise RecNciI were shown in figure 1.

### Non-coding SNVs are also detected, and false positives can be identified

We reviewed all other SNV calls, and noted several known SNPs present in the heterozygous or homozygous state, with quality scores also >500 (Table S6). We also noted seven SNVs that were reported in one or (usually) several samples with low quality scores (all but one <200), all but one intronic (Table S6). These were always transitions (G>A, A>G, or C>T). These base changes were identified as common errors by NanoOK (occurring in 13.31%,12.66%, and 11.95% on average of the relevant base respectively). Furthermore, review of these positions on IGV in all samples revealed a high percentage of reads with the aberrant base, including those where the SNV was not called (11-31%; Figure S4). We concluded that these were false positives. Some were shared by Graphmap and NGMLR alignments from the same sample. Overall, however, the NGMLR alignments had significantly fewer false positives, mostly due to one SNP that was always called in Graphmap samples, but never in NGMLR (mean per sample 2.2 with Graphmap, and 1.2 with NGMLR; paired t-test p=0.0038). For the mean quality of NGMLR false positives, there was no correlation with coverage (r=0.07, p=0.91, for the seven samples which had at least one).

### Structural variant detection and mutation phasing provides additional relevant information

Sniffles and Nanopolish both reported a 55-bp exonic deletion in S5 in the NGMLR-alignment only, clearly visible on IGV in this alignment (Figure S5). This sample had been previously designated “RecNciI” based on the presence of three pseudogene derived missense changes which comprise this genotype. Our detection of the additional missense change p.D448H, and the 55-bp deletion, both of which may coexist with the “RecNciI” mutations, would change the classification to a “c.1263del+RecTL allele”, indicating a different site of recombination with the pseudogene than RecNciI.[8] Detecting this deletion can be difficult with Illumina targeted sequencing.[9] No other structural variants were reported.

We next phased all variants using Whatshap (Table S7). We verified that the four coding SNVs and the deletion in S5 were *in cis*, as well as five rare intronic SNPs already detected in the original analysis (Figure 3). We confirmed compound heterozygosity in two GD cases, S7, heterozygous for p.N409S and p.L105P, and S15, heterozygous for p.R502C and p.R535C. We noted a haplotype comprising 8 SNPs over 6.7 kb. This corresponds to the previously reported Pv1.1^+/−^ haplotype,[25] later extended to a 70-kb haplotype designated 111.[26] One sample was homozygous and two heterozygous for Pv1.1^+^ (Table S7). p.N409S (N370S) was always on the Pv1.1^−^ background, as expected.[8] The p.L483P (L444P) mutation was on the Pv1.1^−^ haplotype in two individuals and the Pv1.1^+^ in one, consistent with the reported lack of founder effect.[8] p.L105P and the complex recombinant allele were on a Pv1.1^−^ haplotype, and p.R502C and p.R535C on Pv1.1^+^.

**Figure 3.**
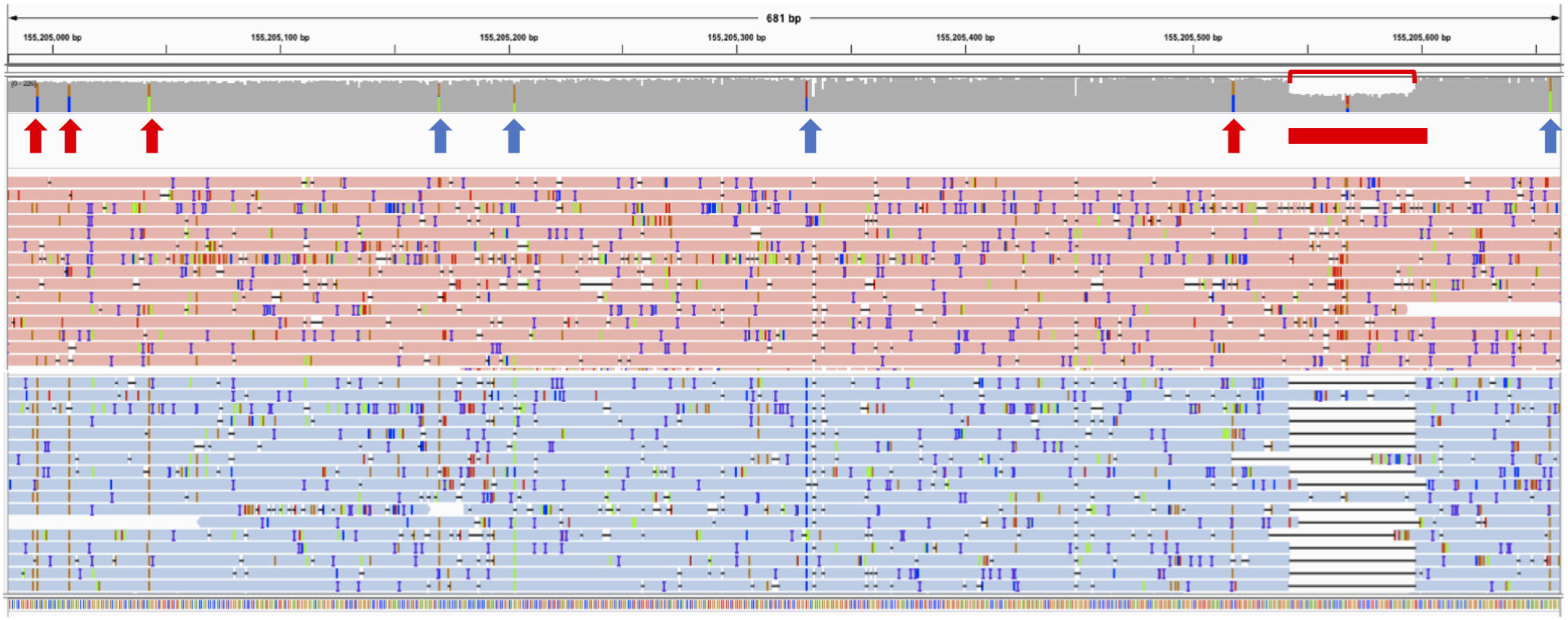
Detection and phasing of a 55-base pair exonic deletion in S5. The coverage track, with eight SNVs highlighted, and a selection of reads are shown, over exons 9 and 10 (chr1:155,204,981-155,205,661; NGMLR alignment). The deletion is clearly visible as a drop in coverage (red bracket). Reads are grouped and coloured by haplotype for these variants, which are all on the blue-coloured reads. The arrows point to the SNVs (red= coding, blue= non-coding) and the red box to the deletion.

## Discussion

We have sequenced a long-range *GBA* amplicon, covering all coding exons and introns, using the Oxford Nanopore Technologies MinION. We first validated our approach on brain DNA samples, using the early R7.3 and R9 chemistry. We then used the newer R9.4 chemistry, as used in human whole genome sequencing,[20] to analyse samples mostly known to carry biallelic or heterozygous mutations, in a blinded fashion. We confirmed common mutations in six samples (p.N409S, p.L483P), differentiating p.N409S homozygosity and heterozygosity. We also detected other mutations in two GD patients, and confirmed compound heterozygosity by phasing mutations where relevant. We further characterised the complex allele previously reported as RecNciI in one PD patient, finding another missense change and a 55-base pair deletion *in cis*, both reported with it before.[8]

Recent years have seen the introduction of single-molecule sequencing in real time by Oxford Nanopore and PacBio which can easily generate long reads of several kb,[17] and in the case of the Nanopore up to hundreds of kb.[20] Using long reads has several advantages, despite the lower accuracy at the base level,[17] some of which were evident here. The challenge of aligning short reads to regions with high homology is often not fully appreciated,[14] with false negatives in *GBA* targeted Illumina sequencing when the whole genome was used as a reference.[9] We observed minimal alignment to the pseudogene. We also detected a coding 55-bp deletion, which can be missed by Illumina data, [9] and intronic SNPs, an understudied area in *GBA* and other lysosomal disorders.[9] Finally, the long reads allowed the phasing of mutations, enabling a haplotype-resolved personalized assessment. This helps overcome the frequent problem of phasing, which may require analysis of relatives.[27]

The nanopore chemistry, and bioinformatic tools available, have evolved considerably during the time in which this work was performed. We compared two aligners (Graphmap and the recently developed NGMLR), both of which gave negligible alignment to the pseudogene. NGMLR allowed detection of the 55-bp deletion, and halved the number of false positives. We thus recommend using NanoOK for quality control, NGMLR for alignment, Nanopolish for SNV calling, and Sniffles for structural variant calling. Nanopolish has been designed for SNV calling by correcting accuracy problems arising in nanopore default basecalling by reanalysing the raw signal data.[20] Nanopolish variant calling option uses a likelihood-based method to generate haplotypes that serve as the reference sequence for the target region.[19] It has been instrumental in projects ranging from Ebola virus[19] to human genome sequencing.[20] A cut-off quality score of 320 in our work differentiated all true and false positives, although the quality score of true positives may partly depend on coverage, and calls with scores ~200-400 would require careful review, and possible Sanger analysis. Although even 120x coverage allowed detection of a p.N409S homozygote, we recommend 300x or more to ensure accurate determination of zygosity. In a human genome sequencing study with SNP analysis of chromosome 20, coverage of only 30x remarkably allowed SNP calling with accuracy ~95% against annotated variants, but zygosity was not always correctly determined.[20]

We were able to identify and filter false positives, based on (1) the low quality score on Nanopolish, (2) the high % of these changes occurring as errors based on NanoOK, and (3) the significant percentage of aberrant bases at the same positions in all samples, even where not called as mutations. Notably, they were always transitions, which were also the main errors in whole genome sequencing using the MinION.[20] Current limitations include the inability to accurately resolve homopolymers and detect small insertions and deletions (indels),[20, 28] and we did not attempt to do this, filtering any indel calls <5 bases. Sniffles can detect insertions and deletions, as demonstrated here, as well as complex structural variants.[28] Based on the rapid developments in the chemistry and bioinformatics, we expect calling of small indels and further reduction of false positive SNV calls in the very near future.

As treatments are now available, neonatal screening for lysosomal storage diseases is becoming commoner,[29] including in some cases Gaucher.[30, 31] This relies on biochemical activity, often by blood-spot screening,[32] with several false positives in Gaucher, possibly due to carrier status.[31] Genetic confirmation is ultimately required, so a rapid and cost-effective method would be useful in this setting. The advantages of the MinION include the very low capital cost, space requirements, and turnaround time of the analysis. The cost per sample is likely to compare favourably with Sanger and Illumina sequencing in all settings, especially taking into account the ability to phase variants and detect structural variants in this complex region. Current R9.4 flow cells yields are at least 5 Gb of sequence, and often much more. For our 8.9 kb amplicon, 96 samples, which can be multiplexed on a single flow cell, would therefore achieve a mean coverage >1,000x, even if less than a fifth of the reads aligned successfully.

## Conclusion

Oxford Nanopore is a versatile single-molecule real-time sequencing technology that has been used in several innovative applications, from detection of Ebola to proof-of-principle human whole genome sequencing. Here we demonstrate that the MinION can detect and phase pathogenic variants in *GBA*, and intronic SNPs that would not be detected by Sanger sequencing of exons. The rapid evolution of specific bioinformatic methods, and the improvements in accuracy and data yield, combined with the minimal footprint and capital investment, make the MinION a suitable platform for long-read sequencing of difficult genes such as *GBA*, both in the diagnostic and research environments.

## Methods

### Overview, DNA extraction and PCR

The aim of this study was to design and validate a protocol for *GBA* analysis using a long amplicon on the MinION. Samples used in this study were derived from 17 individuals (Table S1). We used brain DNA from eight PD patients, one MSA, and one control, including two samples from brains of PD patients known to carry *GBA* mutations RecNciI and p.L483P (L444P).[16] Brain samples were provided by Queen Square and Parkinson’s UK brain banks. We also used samples from saliva of seven living individuals, six of whom had previously been found to have at least one mutation in GBA, although these results were not known to MLS and CP, who performed the SNV analyses, until it had been completed. All individuals had given written informed consent. Ethics approval was provided by the National Research Ethics Service London – Queen Square ethics committee for living individuals, and the National Research Ethics Service London – Hampstead ethics committee for the deceased, with additional permission for study of brains from the research tissue banks provided by the UK National Research Ethics Service (07/MRE09/72). DNA was isolated from brain using Phenol-Chloroform,[33] and from saliva using Oragene-DNA kit.

We amplified an 8.9-kb *GBA* sequence, which covered all coding exons, the introns between them, and part of the 3’ UTR region (chr1: 155202296-155211206; Figure S1). We customised previously reported primers[34] to carry Oxford Nanopore adapters and barcodes for multiplexing. Primer sequences were npGBA-F: 5’- TTTCTGTTGGTGCTGATATTGCTCCTAAAGTTGTCACCCATACATG-3’ and npcMTX1: 5’- ACTTGCCTGTCGCTCTATCTTCCCAACCTTTCTTCCTTCTTCTCAA-3’.

Two DNA polymerases with appropriate optimised PCR conditions were used to amplify the *GBA* target region (Table S2): Expand Long Template PCR (Roche) and Kapa Hi-Fi polymerase (Kapa Biosystems). Amplicons were purified by Qiaquick PCR purification kit (Qiagen) and DNA concentration was measured by Qubit.

### Barcoding, Library Preparation and Sequencing

For sample multiplexing, a barcoding step was carried out after generating the *GBA* amplicons with PCR Barcoding Kit I (Oxford Nanopore). Up to 12 samples were pooled in each case for library preparation according to the manufacturer amplicon sequencing protocol, starting with 1 µg of DNA and 1% DNA CS spike-in for the dA-tailing step, followed by purification using AMPure beads. Nanopore adapters were ligated to the end-prepped DNA, using the NEB blunt/TA ligase master mix recommended by the manufacturer. Flow cell priming was performed according to the requirements of each flow cell version. We first used R7.3 and R9 flow cells with 2D reads, where a molecule passes through the pore in both directions. After recent technical advances, we used 1D reads from a R9.4 flow cell.

### Bioinformatic analysis

MinKNOW versions 0.51.1.62 and later were used for data acquisition and run monitoring. Metrichor versions v2.38.1033 - v2.40.17 were used for basecalling, de-multiplexing and fast5 file generation. The software divides reads into “pass” and “fail”, and only “pass” reads were analysed. We used NanoOK (version 1.25)[35] to obtain a wide range of quality control metrics, with Graphmap (version 0.3.0)[21] alignment, using the precise region targeted as reference. We first converted fast5 files to fastq using NanoOK, or Poretools (version 0.6.0)[36] with a 2-kb size cut-off. NanoOK output included the N50 (the size at which reads of the same or greater length contain 50% of the bases sequenced), the commonest erroneous substitutions, and overall error estimates, notably the aligned base identity excluding indels (ABID), and identical bases per 100 aligned bases including indels (IBAB). We aligned reads to the human genome (hg19) for detailed study and variant calling using GraphMap or NGMLR (version 0.2.6)[28], both specifically developed for long reads. Samtools (version 1.3.1) were used where required to merge, sort and index bam files. Coverage was calculated using bedtools (version 2.25.0) coverageBed. Data were viewed on IGV.

We used Nanopolish (versions 0.6-dev and 0.8.4)[19] to call variants over our target region. Nanopolish was specially developed to improve accuracy by reanalysis of raw signals after alignment, and used in a recent whole genome study.[20] It relies on a hidden Markov model which calculates the probability of the MinION data at the signal-level for a given proposed sequence.[37] We called variants setting ploidy to 2, and invoked the “fix homopolymers” option. When using Nanopolish 0.8.4, we had to use Albacore (version 2.1.3, Oxford Nanopore) to re-generate fastq files for analysis. We filtered any indel calls smaller than 5 bases, due to the known problem of Nanopore in calling these, especially in homopolymer regions.[20, 28] We reviewed the variant quality of all calls and visualised them on IGV. We used WhatsHap, designed for long reads,[38] to phase all true variants, and tag bam files for visualisation. We used Sniffles, another tool designed specifically for such data, to call structural variants.[28]

All nomenclature is based on the Human Genome Variation Society guidelines,[39] using reference sequence NM_000157.3. The traditional numbering for *GBA* missense mutations, which omits the first 39 amino acids, is given in brackets to ensure easy comparability with previous literature. SNVs were annotated using ANNOVAR,[40] and viewed on www.varsome.com, which provides data from dbSNP, gnomAD [41] genomes and exomes where available, and other useful metrics.

### Statistical analysis

This was performed using Graphpad Prism v.6.0 (Graphpad, CA, USA) using paired t-test and Spearman correlation analysis as indicated.

## Declarations

### Ethics approval and consent to participate

All individuals had given written informed consent. Ethics approval was provided by the National Research Ethics Service London – Queen Square ethics committee for living individuals (15/LO/1155), and the National Research Ethics Service London – Hampstead ethics committee for the deceased (10/H0720/21), with additional permission for study of brains from the research tissue banks provided by the UK National Research Ethics Service (07/MRE09/72).

#### Consent for publication

Not applicable as no identifiable data included.

#### Availability of data and materials

The datasets generated and/or analysed during the current study are not publicly available due to consent and ethics restrictions, but are available from the corresponding author on reasonable request.

#### Funding

Melissa Leija-Salazar is funded by CONACYT. FJS was supported by NHGRI grant UM1 HG008898. Additional funding was received by the Michael J Fox Foundation for Parkinson’s research. The funders had no role in the design of the study and collection, analysis, and interpretation of data.

### Authors’ contributions

CP conceived, designed, and supervised the study, performed bioinformatic analysis (Whatshap and NGMLR), and drafted the final version of the manuscript. MLS made substantial contributions to the acquisition of laboratory data, performed the majority of the bioinformatic analysis and interpretation of data, and prepared the first draft. FJS performed bioinformatic analysis (Sniffles) and contributed substantially to the interpretation of all bioinformatic analyses, and advancing the manuscript. KM made substantial contributions to the acquisition of laboratory data. SM, MT, AD, RS, DH, and AHVS made substantial contributions to the study design, and interpretation of the findings. MA performed and interpreted Graphmap alignment of early data. All authors critically revised the manuscript, and read and approved the final version.

## Acknowledgements

We are grateful to the Queen Square and Parkinson’s UK Brain Banks, and to all individual who donated their brains or DNA samples to research. The Queen Square Brain Bank is supported by the Reta Lila Weston Institute for Neurological Studies and the Medical Research Council UK. The Parkinson’s UK Tissue Bank is funded by Parkinson’s UK, a charity registered in England and Wales (258197) and in Scotland (SC037554). We are grateful to Atul Mehta and Sarah Cable for help recruiting participants. We thank Richard Leggett for support of NanoOK, and Jared Simpson for support of Nanopolish.

