## Supplementary Materials for "Detection of *GBA* missense mutations and other variants using the Oxford Nanopore MinION" (from another version of this preprint: version 2, 2018-03-24)

**Supplementary table 1. All samples tested in this study, with diagnosis and DNA source.** Note that two additional brain samples, coded 9 and 10, only gave a few correctly aligned reads, and were not analysed further.

| Sample | Diagnosis | DNA source | age | gender |
|---|---|---|---|---|
| S1 | PD | cerebellum | 73 | m |
| S2 | Control | substantia nigra | 94 | f |
| S3 | PD | substantia nigra | 64 | m |
| S4 | PD | substantia nigra | 79 | f |
| S5 | PD | substantia nigra | 65 | f |
| S6 | MSA | substantia nigra | 56 | m |
| S7 | PD | substantia nigra | 72 | f |
| S8 | PD | substantia nigra | 60 | f |
| S11 | PD | substantia nigra | 75 | m |
| S12 | GD | substantia nigra | 60 | f |
| S13 | healthy carrier | substantia nigra | 32 | f |
| S14 | GD | saliva | 53 | m |
| S15 | GD | saliva | 50 | m |
| S16 | PD | saliva | 80 | m |
| S17 | GD | saliva | 42 | n |
| S18 | PD | substantia nigra | | |
| S19 | healthy carrier | saliva | 71 | m |

**Supplementary Table 2**. PCR conditions

| | Conditions | Protocol | |
|---|---|---|---|
| | | **Roche** | **Kapa** |
| **Reaction** | DNA input (ng) | 250-400 ng | 500 ng |
| | Buffer | Buffer 2 | 2x Kapa HiFi HotStart Ready Mix |
| | Primers | npGBA-F & npcMTX1 | npGBA-F & npcMTX1 |
| | Enzyme | Expand Long template | Kapa HiFi |
| | Final reaction volume (µL) | 50 | 50 |
| **Programme** | Cycles | 35 | 28 |
| | Initial denaturation | 94°C/2min | 95°C/3min |
| | Denaturation | 94°C/10sec | 98°C/20sec |
| | Annealing | 57°C/30sec | 60.9°C/15sec |
| | Extension | 68°C /7min (increasing 20 sec each cycle from 11 onwards) | 72°C/9min |
| | Final extension | 68°C/9min | 72°C/9min |

**Supplementary Table S3. Summary of all runs.**

The sequencing conditions, read metrics, PCR protocol used, and mean aligned base identity assessment (ABID) by NanoOK are shown for each sequencing run. Barcoding was used in all runs expect the first one. Bam files from the same samples in runs 1-5 were merged prior to variant calling. (*) This flow cell was reused after washing, leading to very low yield.

| Run parameters | Run number | | | | | |
|---|---|---|---|---|---|---|
| | **1** | **2** | **3** | **4** | **5** | **6** |
| Type of reads | 2D | | | | | 1D |
| Flow cell version | R7.3 | R7.3 | R7.3* | R7.3 | R9 | R9.4 |
| Run time (hours) | 2 | 48 | 46 | 48 | 48 | 48 |
| PCR protocol | Roche | Roche | Roche | Mostly Kapa | Mostly Kapa | Kapa |
| Samples analysed | 1 | 9 | 3 | 7 | 10 | 10 |
| Mean ABID (%) ± SD | 94.62 | 94.96 ±0.19 | 94.84 ±0.29 | 94.78 ±0.76 | 96.45 ±1.55 | 93.56 ±1.21 |

**Supplementary table 4. Detailed data for validation runs 1-5. Samples tested, PCR protocol used, NanoOK metrics, alignment compariso and total numbers of SNPs called**

NanoOK quality assessment after alignment to the target sequence (chr1:155202292-155211206) is shown. The coverage of the gene and pseudogene (equivalent region over *MTXN-GBAP* chr1:155181579-155189601) after Graphmap alignment to the whole genome and mergi bam files from different runs is also given. Total SNPs includes the coding ones in S5 and S8. Note that S7 was included in runs 4 and 5 as separately barcoded samples, amplified with different PCR enzymes. There was a possible minimal improvement in ABID with Kapa in both cases (1% and 0.1%).

| | **Experiment** | | **NanoOK analysis** | | | | **Coverage** | | **Total non-coding SNP** |
|---|---|---|---|---|---|---|---|---|---|
| **Sample** | **Run ID** | **PCR protocol** | **N50** | **Reads aligned (%)** | **ABID (%)** | **IBAB (%)** | **gene** | **pseudogene** | |
| S1 | 1 | Roche | 8783 | 94.6 | 95.0 | 85.5 | 579 | 11 | 12 |
| | 2 | Roche | 8734 | 37.5 | 95.4 | 85.8 | | | |
| | 3 | Roche | 8775 | 40.4 | 95.4 | 85.9 | | | |
| | 4 | Kapa | 8464 | 100.0 | 95.7 | 85.4 | | | |
| | 5 | Kapa | 8454 | 33.1 | 97.4 | 89.9 | | | |
| S2 | 2 | Roche | 8737 | 44.0 | 94.8 | 84.5 | 63 | 0 | 2 |
| S3 | 2 | Roche | 8714 | 48.6 | 95.0 | 85.0 | 359 | 6 | 10 |
| | 3 | Roche | 8752 | 46.0 | 94.8 | 84.7 | | | |
| | 4 | Kapa | 8685 | 86.2 | 95.7 | 86.2 | | | |

| | | | | | | | | | |
|---|---|---|---|---|---|---|---|---|---|
| | 5 | Kapa | 8486 | 74.9 | 97.3 | 89.8 | | | |
| S4 | 2 | Roche | 8687 | 33.1 | 95.1 | 84.9 | 79 | 1 | 2 |
| | 5 | Kapa | 524 | 4.9 | 93.5 | 82.9 | | | |
| S5 | 2 | Roche | 8725 | 36.1 | 95.0 | 84.6 | 235 | 2 | 8 |
| | 3 | Roche | 8824 | 41.7 | 94.9 | 84.8 | | | |
| | 4 | Roche | 8632 | 87.5 | 95.3 | 85.5 | | | |
| | 5 | Roche | 8663 | 86.5 | 97.4 | 90.1 | | | |
| S6 | 2 | Roche | 8740 | 39.4 | 89.5 | 84.7 | 113 | 2 | 7 |
| | 5 | Kapa | 4215 | 8.9 | 98.1 | 91.6 | | | |
| S7 | 2 | Roche | 8722 | 31.2 | 94.7 | 84.3 | 291 | 0 | 12 |
| | 4 | Roche/Kapa | 8595/8603 | 100/100 | 94.7/95.7 | 84.7/85.9 | | | |
| | 5 | Roche/Kapa | 8660/8572 | 87.9/74.1 | 97.3/97.4 | 89.9/90.1 | | | |
| S8 | 2 | Roche | 8726 | 64.7 | 95.1 | 85.0 | 1095 | 9 | 2 |
| | 4 | Kapa | 8649 | 98.2 | 95.7 | 86.0 | | | |
| | 5 | Kapa | 8618 | 88.8 | 97.5 | 90.2 | | | |
| S11 | 2 | Roche | 8743 | 38.6 | 94.9 | 84.7 | 70 | 1 | 6 |
| | 5 | Kapa | 761 | 4.9 | 96.1 | 88.1 | | | |
| | | | | | | **Mean** | **320.4** | **3.6** | **6.8** |

**Supplementary Table S5. R9.4 (run 6) NanoOK metrics, and coverage of gene and pseudogene by each aligner.**

| Sample | NanoOK metrics | | | | Coverage by each aligner | | | |
|---|---|---|---|---|---|---|---|---|
| | | | | | Graphmap | | NGMLR | |
| | N50 | % reads aligned | ABID (%) | IBAB (%) | GBA | pseudo-gene | GBA | pseudo-gene |
| S5 | 8851 | 73.5 | 94.6 | 84.3 | 250.7 | 3.4 | 234.8 | 11.7 |
| S8 | 4052 | 32.3 | 92.5 | 81.3 | 435.2 | 4.8 | 427.2 | 9.0 |
| S12 | 8440 | 40.5 | 94.6 | 84.5 | 270.5 | 2.2 | 264.0 | 4.3 |
| S13 | 8737 | 55.3 | 94.5 | 84.3 | 499.1 | 4.6 | 496.6 | 7.1 |
| S14 | 3976 | 31.3 | 92.4 | 81.3 | 121.8 | 3.2 | 119.1 | 2.9 |
| S15 | 7465 | 45.3 | 93.6 | 82.9 | 400.3 | 1.7 | 392.2 | 7.0 |
| S16 | 3634 | 27.5 | 91.4 | 79.9 | 331.9 | 0.7 | 322.7 | 5.6 |
| S17 | 4543 | 38.0 | 93.6 | 82.9 | 208.0 | 2.2 | 205.1 | 3.3 |
| S18 | 8888 | 64.6 | 94.8 | 84.9 | 285.6 | 0.5 | 280.2 | 3.4 |
| S19 | 3865 | 24 | 92.0 | 79.7 | 415.9 | 1.2 | 409.3 | 7.4 |
| mean | | | | | 322 | 2.5 | 315 | 6.2 |
| % of coverage of pseudogene to gene | | | | | 0.78 | | 1.97 | |

**Commented [MOU2]:** As before- remove these?

**Supplementary table S6. Other SNVs, including false positives, reported in run 6 after NGMLR alignment.**

Details are shown for true and false positives separately. For known SNPs detected, the position, ID, and population allelic frequency are given. For each SNV called in a sample, the Nanopolish quality score is given. The mean quality score for all true positive heterozygotes, and false positives, in each sample is shown (in brackets if there was only one call). False positive number 3 was reported in all Graphmap-aligned samples, but not in NGMLR. The total false positive calls per sample with each aligner is shown at the end.

| | | Sample | S5 | S8 | S12 | S13 | S14 | S15 | S16 | S17 | S18 | S19 |
|---|---|---|---|---|---|---|---|---|---|---|---|---|
| **True positive non-coding SNPs** | | | | | | | | | | | | |
| | Position (ID if applicable) | Base change | | | | | | | | | | |
| 1 | 155202689 rs10796940 | A>C | | | | 1161.7 | | 2307.4 | | | | 881.7 |
| 2 | 155203060 rs11264344 | G>A | | | | 754 | | 1569.6 | | | | 606.2 |
| 3 | 155204694 rs708606 | C>T | 546.9 | | | | | | | | | |
| 4 | 155205170 rs2974924 | A>G | 533 | | | | | | | | | |
| 5 | 155205203 rs426516 | G>A | 551.4 | | | | | | | | | |
| 6 | 155205331 rs1800473 | T>C | 677.1 | | | | | | | | | |
| 7 | 155205657 rs111417507 | A>G | 1458.5 | | | | | | | | | |
| 8 | 155205669 rs3115534 | G>T | 1458.5 | | | | | | | | | |
| 9 | 155205709 rs548435731 | A>G | 822.2 | | | | | | | | | |
| 10 | 155206341 rs9628662 | T>G | | | | 1478.9 | | 2664.4 | | | | 1433.2 |

| | | | | | | | | | | | | |
|---|---|---|---|---|---|---|---|---|---|---|---|---|
| 11 | 155207030<br>rs72704130 | G>A | | | | | | 1211 | | | | |
| 12 | 155207549<br>rs762488 | T>C | | | | 721.6 | | 1486.9 | | | | |
| 13 | 155207626<br>rs2009578 | G>A | | | | 1392.2 | | 2624.3 | | | | 684.2 |
| 13 | 155208183<br>rs2974923 | T>C | | | | 1011.9 | | 1745.2 | | | | 834.7 |
| 14 | 155208647 | T>C | 1353.3 | 2303.3 | 1479.3 | 2727 | 588.2 | | 1776.7 | 1079.2 | | 2228.6 |
| 15 | rs7416991 | T>G | | | | | | 2268.3 | | | | |
| 16 | 155209360<br>rs2075569 | C>T | | | | 916.5 | | 1614.3 | | | | 707.2 |
| Mean of **heterozygote** calls<br>(including coding SNVs from table 1) | | | 775 | (630.3) | | 1039 | | 1126 | | 382 | | 839 |
| **False positives** | | | | | | | | | | | | |
| 1 | 155203446 | G>A | | | | | 105.3 | | | | | |
| 2 | 155203545 | C>T | | | | | | 103.4 | | | | |
| 3 | 155204190 | G>A | | | | | | | | | | |
| 4 | 155204802 | A>G | 99.4 | | | | | | 127.1 | | | |
| 5 | 155206733 | G>A | | | | | 24.6 | | | | | |
| 6 | 155210127 | C>T | 29.2 | 26.8 | | | | 41.3 | | | | |
| 7 | 155211111 | G>A | 143.6 | | 175.2 | 309.3 | 113.1 | | | | | |
| | | Mean | 91 | (26.8) | (175.2) | (309.3) | 81 | 72 | (127.1) | | | |
| False positive calls by aligner | | Graphmap | 3 | 3 | 2 | 2 | 4 | 2 | 1 | 2 | 2 | 1 |
| | | NGMLR | 3 | 1 | 1 | 1 | 3 | 2 | 1 | 0 | 0 | 0 |

**Supplementary table S7. All SNPs with phasing and population allele frequencies.**

Coding mutations are highlighted in red. The DNA changes labelled with (*) constitute RecNciI when together. The common SNPs that comprise the haplotype Pv1.1 are in yellow. Note that two different SNPs were found at position 155208647, with the combined AF 95%. The overall frequency of the (+) haplotype, calculated as the mean AF of individual SNP comprising it, is ~38.3%. This was previously estimated to constitute 30% of Caucasian and 70% of African and Asian haplotypes.[1]

**Supplementary Figure S1. GBA gene structure.**

Image shows exact region targeted by our primers (chr1: 155202296-155211206).

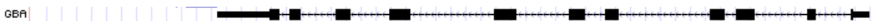


**Supplementary Figure S2. Intronic variant detected in sample S1 by MinION.**

(A) IGV shows the read count for each base at this position, consistent with heterozygosity.

(B) Sanger sequencing confirmed this variant to be real (arrow). Primer used GCGCCATCTTCACTCACTGTAAC

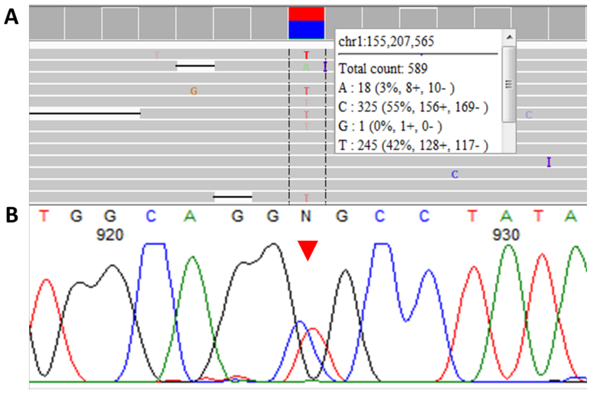

**Supplementary figure S3. False positive in sample S3.** (chr1: g.155210582A>G)

(A) IGV coverage and supporting reads for each base at the variant position demonstrates that it was present in 6 of the 9 samples above 10% AF, including sample S3 (red box).

(B) Sanger sequencing using primer ACTGCCTTGACTCACTCACC. This position (red arrow) has the reference allele.

(C) NanoOK table of common substitutions  shows that A>G is a common sequencing error.

A

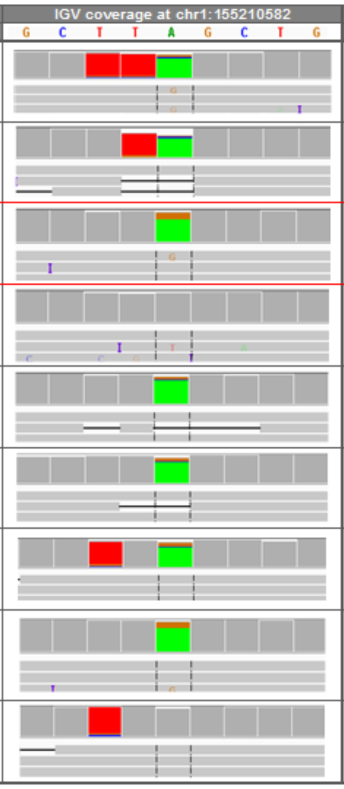


| Sample | % of reads showing each base | | | |
|---|---|---|---|---|
| Ref | A | C | G | T |
| S1 | 83 | 6 | 10 | 1 |
| S2 | 86 | 10 | 4 | 0 |
| S3 | 75 | 0 | 24 | 1 |
| S4 | 92 | 4 | 3 | 1 |
| S5 | 86 | 3 | 10 | 1 |
| S6 | 83 | 5 | 12 | 0 |
| S7 | 81 | 3 | 16 | 0 |
| S8 | 79 | 1 | 20 | 0 |
| S11 | 94 | 1 | 3 | 0 |

B

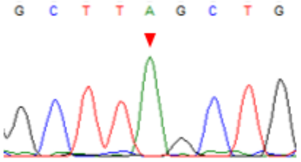


C

| | 2D substituted % | | | |
|---|---|---|---|---|
| | a | c | g | t |
| Reference A | 0.00 | 6.17 | 15.94 | 6.00 |
| Reference C | 4.31 | 0.00 | 3.94 | 10.56 |
| Reference G | 12.99 | 5.17 | 0.00 | 5.34 |
| Reference T | 5.07 | 19.29 | 5.24 | 0.00 |

**Supplementary Figure S4. False positives SNV calls with R9.4 chemistry.**

The numbers correspond to Supplementary Table S8. Sample in which a given SNV was called after Graphmap alignment are denoted by ^, and after NGMLR by *. All the samples from this flow cell are shown, with the number of calls of each base at that position, illustrating the high number of errors at these positions in all or most samples. Note that (3) was always called in Graphmap aligned samples, and never in NGMLR-aligned ones.

**(1)**

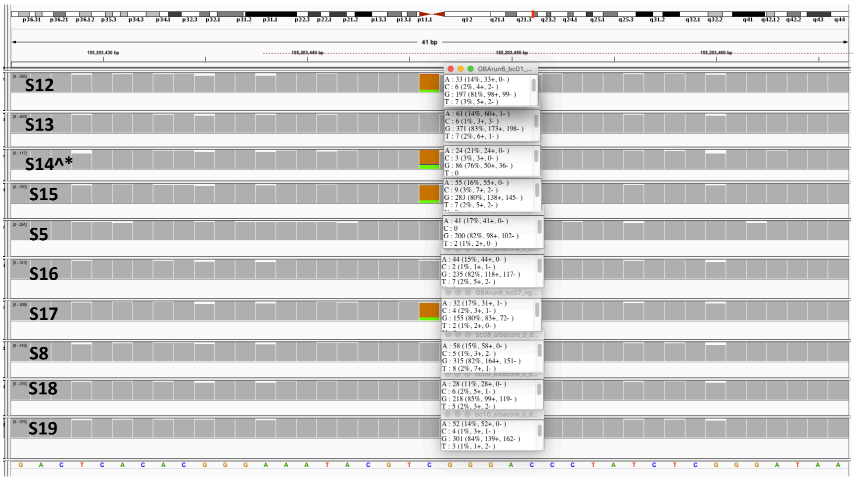


**Commented [LSM3]:** I can help making all figures uniform

(2)

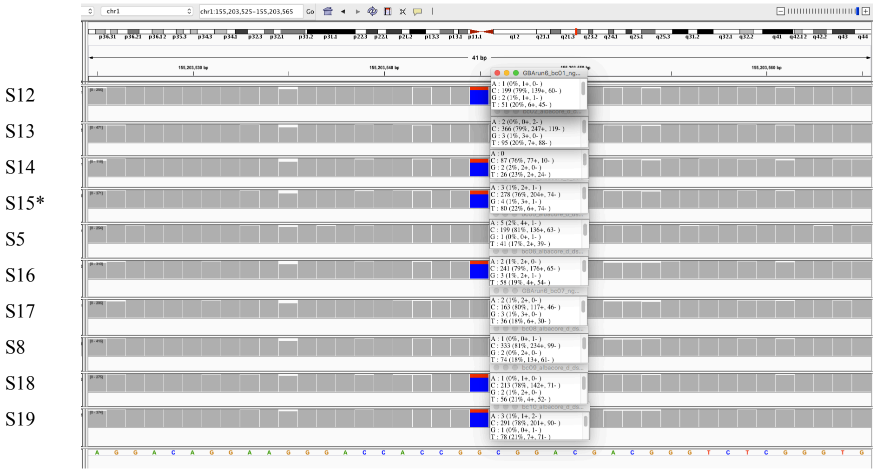
S12
S13
S14
S15*
S5
S16
S17
S8
S18
S19

(3)

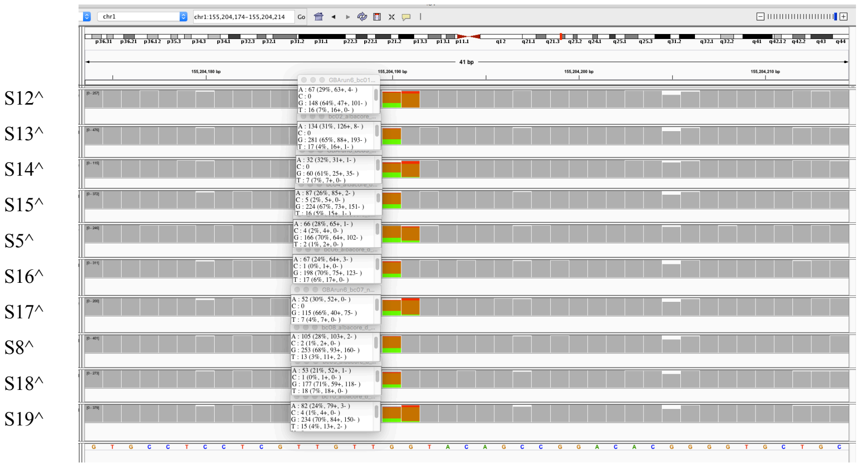
S12^
S13^
S14^
S15^
S5^
S16^
S17^
S8^
S18^
S19^

**(4)**

S12
S13
S14
S15
S5^*
S16^
S17
S8
S18
S19

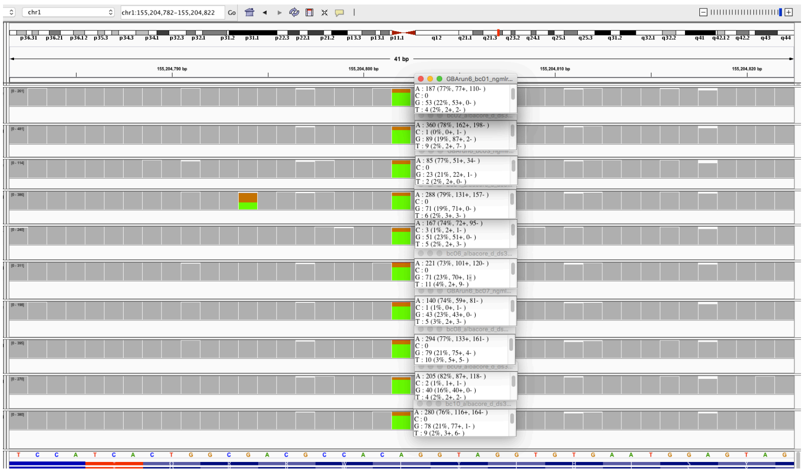

**(5)**

S12
S13
S14^*
S15
S5
S16
S17
S8^
S18
S19

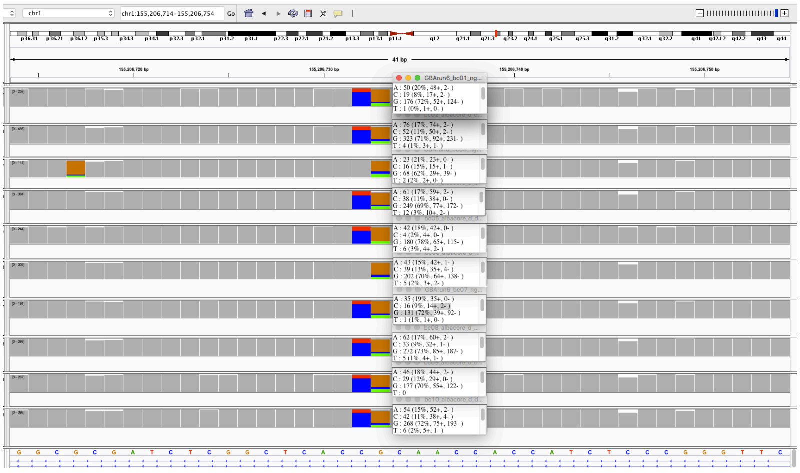

**(6)**

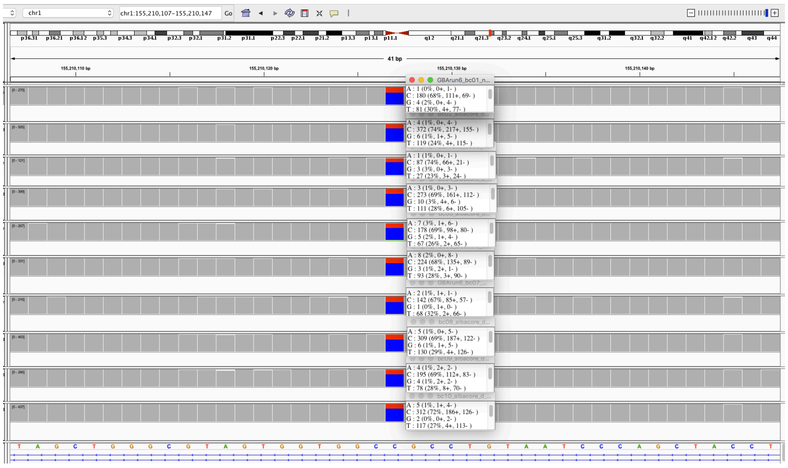

S12
S13
S14
S15^*
S5^*
S16
S17^
S8^*
S18
S19

**(7)**

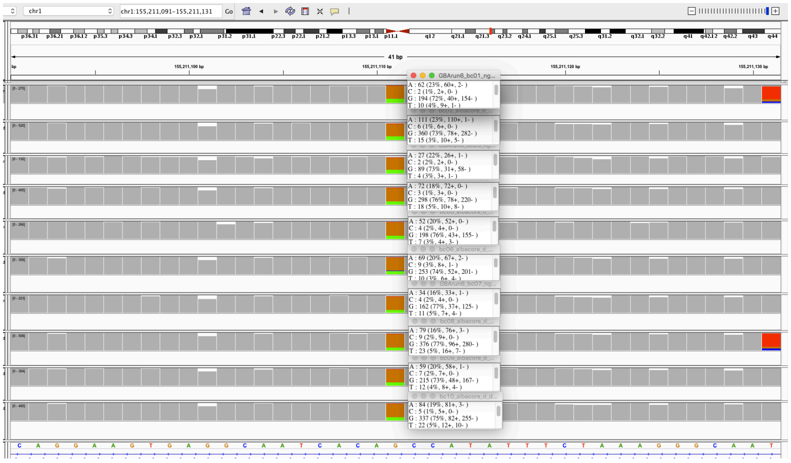

S12^*
S13^*
S14^*
S15
S5*
S16
S17
S8
S18^
S19

**Supplementary Figure S5. Visualisation of 55-base pair exonic deletion in S5 with different aligners.**

Top: NGMLR alignment shows clear drop in coverage, and reads spanning deletion.

Bottom: Graphmap alignment does not clearly reveal deletion.

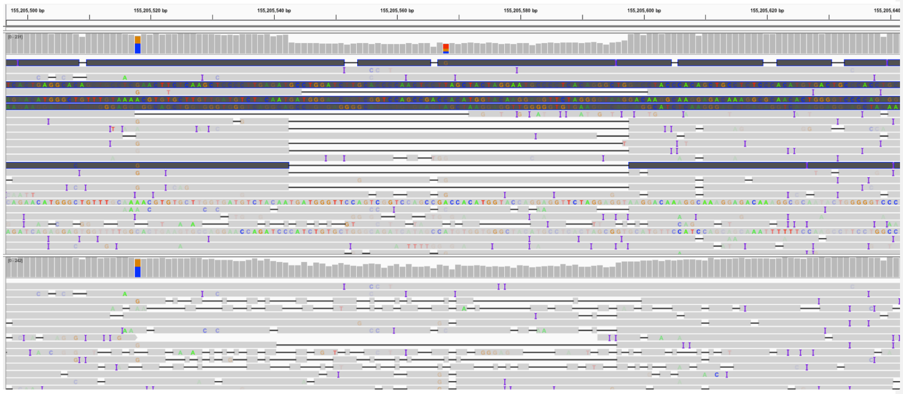
