## Supplementary Materials for "Detection of *GBA* missense mutations and other variants using the Oxford Nanopore MinION" (from another version of this preprint: version 2, 2018-03-24)

| SNP location | ID | ref | alt | DNA change | protein change | intron / exon | AF | S12 | S13 | S14 | S15 | S5 | S16 | S17 | S8 | S18 | S19 |
|---|---|---|---|---|---|---|---|---|---|---|---|---|---|---|---|---|---|
| 155202689 | rs10796940 | A | C | | | 3 in MTXP1 | 0.38830 | | 0 \| 1 | | hom | | | | | | 0 \| 1 |
| 155203060 | rs11264344 | G | A | | | 4 in MTXP1 | 0.38620 | | 0 \| 1 | | hom | | | | | | 0 \| 1 |
| 155204694 | rs708606 | C | T | c.92G>A | | ex11 (UTR) | 0.00219 | | | | | 0 \| 1 | | | | | |
| 155204794 | rs747506979 | G | A | c.1603C>T | R535C | ex11 | 0.00010 | | | | 0 \| 1 | | | | | | |
| 155204987 | rs80356771 | G | A | c.1504C>T | R502C | ex10 | 0.00000 | | | | 1 \| 0 | | | | | | |
| 155204994 | rs1135675 | C | G | c.1497G>C* | V499V | ex10 | 0.00031 | | | | | 0 \| 1 | | | | | |
| 155205008 | rs368060 | C | G | c.1483G>C* | A495P | ex11 | 0.00014 | | | | | 0 \| 1 | | | | | |
| 155205043 | rs421016 | A | G | c.1448T>C* | L483P | ex10 | 0.00131 | | 0 \| 1 | | | 0 \| 1 | | | het | | 1 \| 0 |
| 155205170 | rs2974924 | A | G | c.1389-68T>C | | intron 9 | 0.01053 | | | | | 0 \| 1 | | | | | |
| 155205203 | rs426516 | G | A | c.1389-101C>T | | intron 9 | 0.00298 | | | | | 0 \| 1 | | | | | |
| 155205331 | rs1800473 | T | C | c.1388+141A>G | | intron 9 | 0.37950 | | 0 \|1 | | hom | 0 \| 1 | | | | | 0 \|1 |
| 155205518 | rs1064651 | C | G | c.1342G>C | D448H | ex9 | 0.00026 | | | | | 0 \| 1 | | | | | |
| 155205634 | rs76763715 | T | C | c.1226A>G | N409S | ex9 | 0.00165 | hom | | hom | | | | 0 \| 1 | | | |
| 155205657 | rs111417507 | A | G | c.1225-22T>C | | intron 8 | 0.00023 | | | | | 0 \| 1 | | | | | |
| 155205669 | rs3115534 | G | T | c.1225-34C>A | | intron 8 | 0.95200 | hom | hom | hom | hom | 1 \|0 | hom | hom | hom | hom | hom |
| 155205709 | rs548435731 | A | G | c.1225-74T>C | | intron 8 | 0.00045 | | | | | 0 \| 1 | | | | | |
| 155206341 | rs9628662 | T | G | c.1000-81A>C | | intron 7 | 0.38810 | | 0 \| 1 | | hom | | | | | | 0 \| 1 |
| 155207030 | rs72704130 | G | A | c.999+102C>T | | intron 7 | 0.01379 | | | | 1 \| 0 | | | | | | |
| 155207549 | rs762488 | T | C | c.762-180A>G | | intron 6 | 0.38770 | | 0 \| 1 | | hom | | | | | | 0 \| 1 |
| 155207626 | rs2009578 | G | A | c.762-257C>T | | intron 6 | 0.38260 | | 0 \| 1 | | hom | | | | | | 0 \| 1 |
| 155208183 | rs2974923 | T | C | c.589-86A>G | | intron 5 | 0.36060 | | 0 \| 1 | | 1 \| 0 | | | | | | 0 \| 1 |
| 155208647 | rs7416991 | T | C | c.455-206A>G | | intron 4 | 0.56790 | hom | 1 \| 0 | hom | | hom | hom | hom | hom | hom | 1 \| 0 |
| 155208647 | rs7416991 | T | G | c.455-206A>C | | intron 4 | 0.38510 | | 0 \| 1 | | hom | | | | | | 0 \| 1 |
| 155209360 | rs2075569 | C | T | c.454+47G>A | | intron 4 | 0.36590 | | 0 \| 1 | | hom | | | | | | 0 \| 1 |
| 155209547 | | A | G | c.314T>C | L105P | ex4 | 0.00000 | | | | | | | 1 \| 0 | | | |
| | | | | | | | **haplotype** | -/- | -/+ | -/- | +/+ | -/- | -/- | -/- | -/- | -/- | -/+ |
